# Exploring the use of environment-agent-based models for risk assessment of Great Lakes piping plovers

**DOI:** 10.1101/2024.09.25.615010

**Authors:** Brandon P.M. Edwards, Shoshanah Jacobs, Daniel Gillis

## Abstract

The Piping Plover *Charadrius melodus* is an endangered species of shorebird endemic to North America. This species has been the centre of many modelling studies in the last decade. One model type that has been underused in Piping Plover studies is agent-based modelling, which can be used as an accurate risk assessment tool in simulating effects of anthropogenic activities on a given animal species. Recent innovations in ecological modelling have given rise to the environmental agent-based model (enviro-ABM), which efficiently stores information about spatially indexed environmental cells and treat those as agents. This restricts computation to only focus on environmental cells containing the species we are studying, allowing for a more efficient simulation. Using Python, a high-level programming language popular in scientific computing, we develop an enviro-ABM to provide simulations of Piping Plover hatchling growth during a given breeding season. We experiment with increasing levels of human presence and human exclosure size and observe their effects on the growth rate of the simulated Piping Plover hatchlings. Our simulations showed a clear decrease in Piping Plover growth rate as anthropogenic presence increased in the simulated environment. However, when we added a 100 m human exclosure around the nest, the effects of the anthropogenic presence were mitigated at each level. We conclude that an enviro-ABM can be used to assist with conservation and management decisions, with the caveat that the model be constantly updated and informed with results of field studies, especially those pertaining to foraging and energetics of Piping Plovers.

## 1 Introduction

Loss of suitable breeding habitat is a pressing issue across all taxa and environments (e.g. plants: Holzmueller et al. 2006; invertebrates: Flockhart et al. 2015; terrestrial vertebrates: Cushman 2006; avian vertebrates: Dolman & Sutherland 1995). Decline in suitable breeding habitat, largely caused by humans, is a contributing factor in overall abundance declines of species at risk and it is therefore important to protect sensitive breeding habitat from human disturbance, especially for species at risk (Fahrig 1997). Many species at risk can provide functionality to scientists as indicator species, so the study and conservation of these species can offer important insights to the state of the species’ ecosystem (Lawler et al. 2003). With the global population of humans expected to reach close to ten billion by 2050 (United Nations et al. 2017), it is becoming more difficult to balance the need for developed land with the need for maintaining suitable breeding habitat for species at risk. That said, suitable breeding habitat for species at risk is not necessarily an “all-or-nothing” scenario; compromises can exist that may satisfy both the land needs for human activity and the land requirements for breeding species at risk (Fahrig 1997). The challenge arises in determining just how much of a compromise needs to be made for a mutual benefit. There is a need for development of a tool that will model the conflict that exists between land development and conservation of reproductive habitat for species at risk such that it is possible to identify the theoretical point of compromise for the purposes of mutual benefit.

Agent-based models (ABMs) are a type of simulation model that can be used for risk assessment in ecological systems (Grimm et al. 2010). ABMs seek to simulate life processes of one or more animal species, with each animal in the model acting as an agent (Cohen 2014, Carter et al. 2015). In a typical ABM, thousands of individual agents perform autonomous actions based on pre-set rules (Railsback & Grimm 2011). When creating these rules, it is quite common for some of the agents’ behaviour to be unknown in real situations, or there may be uncertainty attached with it. These uncertain behaviours are predicted stochastically in ABMs (Railsback & Grimm 2011). As more rules and parameters are created for agents, the complexity of the ABM increases (Grimm et al. 2005). While the model should be as close to real life as possible, it becomes a challenge to run multiple simulations in a reasonable amount of time (Grimm et al. 2005, Rose et al. 2017).

Rose et al. (2017) proposed an alteration to the traditional agent-based model: an environmental agent-based model. In an environmental agent-based model (enviro-ABM), instead of treating each animal as agents, the habitat (or environment) that the animal species of interest resides in is partitioned into cells, and these cells are treated as agents (Rose et al. 2017). Each of these cells (enviro-agents) stores the total number of individuals for that particular cell. Because the size of the environment is static throughout the simulation, the total number of agents in the simulation remains constant, even though the population of animals could increase or decrease. Additionally, since only a subset of the environment contains animals during a simulation (i.e. only a subset of the agents is active), the number of operations per time step can be restricted to only active agents. Storing the total number of animals in each enviro-agent still allows for stochastic calculations that would be performed on traditional individual agents, but it has the benefit of these animals being sorted by location and age for more efficient calculations (Rose et al. 2017).

This type of model was originally developed and implemented to simulate a large population of Lake Huron Lake Whitefish Coregonus clupeaformis. In the case study, Lake Huron was partitioned into 1 km by 1 km cells, and these cells were treated as the enviro-agents. Each cell stored information related to population, habitat type (based on depth at that point in the lake), spawning, mortality, and movement of individuals. The output of that model was used to make inferences about the impact of differing intensities of anthropogenic activities in the lake environment. Since the agents were based on the environment, it made it easy to add or remove different environmental stressors to observe how the simulated population of Lake Whitefish reacted. An additional benefit of basing the agents directly on the environment was that emergent properties of simulated land-use could be inferred from model runs (i.e. given different scenarios, how do the simulated species use the different parts of the land; Rose et al. 2017). With this in mind, an enviro-ABM is a good candidate to use for simulating species at risk and their interaction with their environment, especially when it comes to energy gain of vulnerable young.

In avian species at risk, protection of habitat is an important step in developing conservation management plans and enviro-ABMs may be of use in assisting with management decisions. In this paper, we describe the implementation of an enviro-ABM for use with the Piping Plover *Charadrius melodus*. We focus on simulating breeding Piping Plovers at Sauble Beach, a popular tourist beach in South Bruce Peninsula, ON. Piping Plovers have regularly bred at Sauble Beach since 2007 following a 30-year absence (Environment Canada 2013). However, because of the beach’s popularity in the summer, Piping Plovers nesting on Sauble Beach face constant direct or indirect human disturbance, whether from humans approaching the plovers, altering the vegetation by raking the beach, or facilitating predation.

The goal of this paper is to explore the use of enviro-ABMs in risk assessment of Piping Plovers breeding at Sauble Beach. It will be necessary to satisfy the following objectives: (1) describe an enviro-ABM for simulating a population of Piping Plover chicks, (2) investigate a set of input parameters to create a base model; that is, a “perfect world” scenario in which there are no anthropogenic disturbances to nesting Piping Plovers and (3) create and evaluate sets of test scenarios with which to compare to the base model. This study only looks at the growth rate in simulated Piping Plover chicks as this can be used as a good indicator of the viability of a Piping Plover chick (Wilcox 1959). This model assumes no predation. The test scenarios mentioned in objective (3) will include (a) investigating how anthropogenic presence affects Piping Plover hatchling growth rate, and (b) investigating how the use of a 100 m human exclosure (that is, a closed off area surrounding the nest) affects Piping Plover hatchling growth rate given differing intensities of anthropogenic presence as in (a). Results from these tests will inform us on the effectiveness of an enviro-ABM to pick up changes in Piping Plover habitat. Recommendations will be made regarding next steps in developing and applying an enviro-ABM toward informing Piping Plover conservation efforts.

## 2 Methods

### 2.1 Environmental Agent-based Model

A typical enviro-ABM consists of three hierarchical layers: physical, biological, and anthropogenic (Rose et al. 2017). The physical layer refers to the abiotic components of the environment in which the animal species of interest resides (e.g. geomorphology, weather, climate, etc.). The biological layer refers to the animal species that the model is simulating and its interactions with the physical layer (e.g. land use, behaviour during certain weather, etc.). The anthropogenic layer describes the human-driven factors that affect the physical layer, biological layer, or both. Enviro-ABMs allow for each layer to be developed independently of each other, allowing for simplicity in designing experiments (as discussed later). As a motivating example, we can classify each of the three layers in the study led by Rose et al. (2017): the physical layer was Lake Huron and its associated bathymetry, the biological layer was the Lake Whitefish and their associated behaviour given the depth or time of year, and one of the anthropogenic layers was harvesting, which, in this case, only affected the biological layer (i.e. the Lake Whitefish abundance).

The model was developed using Python 3.6 (van Rossum 1995). The following subsections describe the development of each the physical, biological, and anthropogenic layers and how they interact with each other. Table 1 breaks down the relevant literature sources from which we based our assumptions for building the model.

**Table 1:** Table of assumptions and associated literature sources for each layer of the model.

| Model Layer | Assumption | Literature Source(s) | Gap in Knowledge |
| --- | --- | --- | --- |
| Physical | Habitat types at Sauble Beach | Canadian Wildlife Service *et al.* (2015), Hamm (2016) | Field studies needed for more specificity in microhabitat |
|  | Foraging quality of habitat | Le Fer *et al.* (2008), Haffner *et al.* (2009), Maslo *et al.* (2011), Cohen (2014) | Site-specific invertebrate studies would be useful for environment-specific foraging quality |
| Biological | Nest creation in sandy beach habitat type | Cairns (1982) |  |
|  | At least 200 m between nests | Cairns (1982) |  |
|  | Clutch size of 4 per nest | Cairns (1982) |  |
|  | Initial piping plover weight (mean 6 grams) | Wilcox (1959), Cairns (1982), Elliott-Smith and Haig (2004), |  |
|  | Random walk (foraging) | Haffner *et al.* (2009), Maslo *et al.* (2011), Cohen (2014), Hamm (2016), | Site-specific observational studies of habitat use would be more beneficial in informing model |
|  | Growth curve parameter estimates | Martens and Goossen (2008) | Growth curve only obtained from one study site; site-specific growth curves would be useful |
|  | No foraging at night | Staine and Burger (1994) | More observational studies needed for night foraging frequency |
| Anthropogenic | Piping plover alert distance | Gulbransen *et al.* (2006), Cohen (2014) |  |
|  | Forage at reduced rate with humans nearby | Burger (1992) |  |

### 2.2 Physical Layer

In this study, the physical layer was a 5366 m × 2040 m area of Sauble Beach, Ontario. Using the polygon tool in ArcMap 10.5, a map of Sauble Beach was digitized into 6 key habitat types: open water, intertidal, open beach, stream, dunes, and urban (Cohen 2014, Canadian Wildlife Service 2015). These habitat types play a role in how the Piping Plover moves around its environment, as each habitat type will have an associated “foraging quality” with it (i.e. open beach habitat type will have better foraging quality than open water habitat type; Maslo et al. 2011). The digitized map was exported as a TIFF file from ArcMap. Georeferencing data were stored in the TIFF file, and so each pixel of the TIFF image was able to be mapped to a 1 m × 1 m plot of land that could be labeled with its specific habitat type. This gives a 5366 × 2040 matrix of habitat types corresponding to 1 m × 1 m plots of land in Sauble Beach (Figure 1).

**Figure 1:**
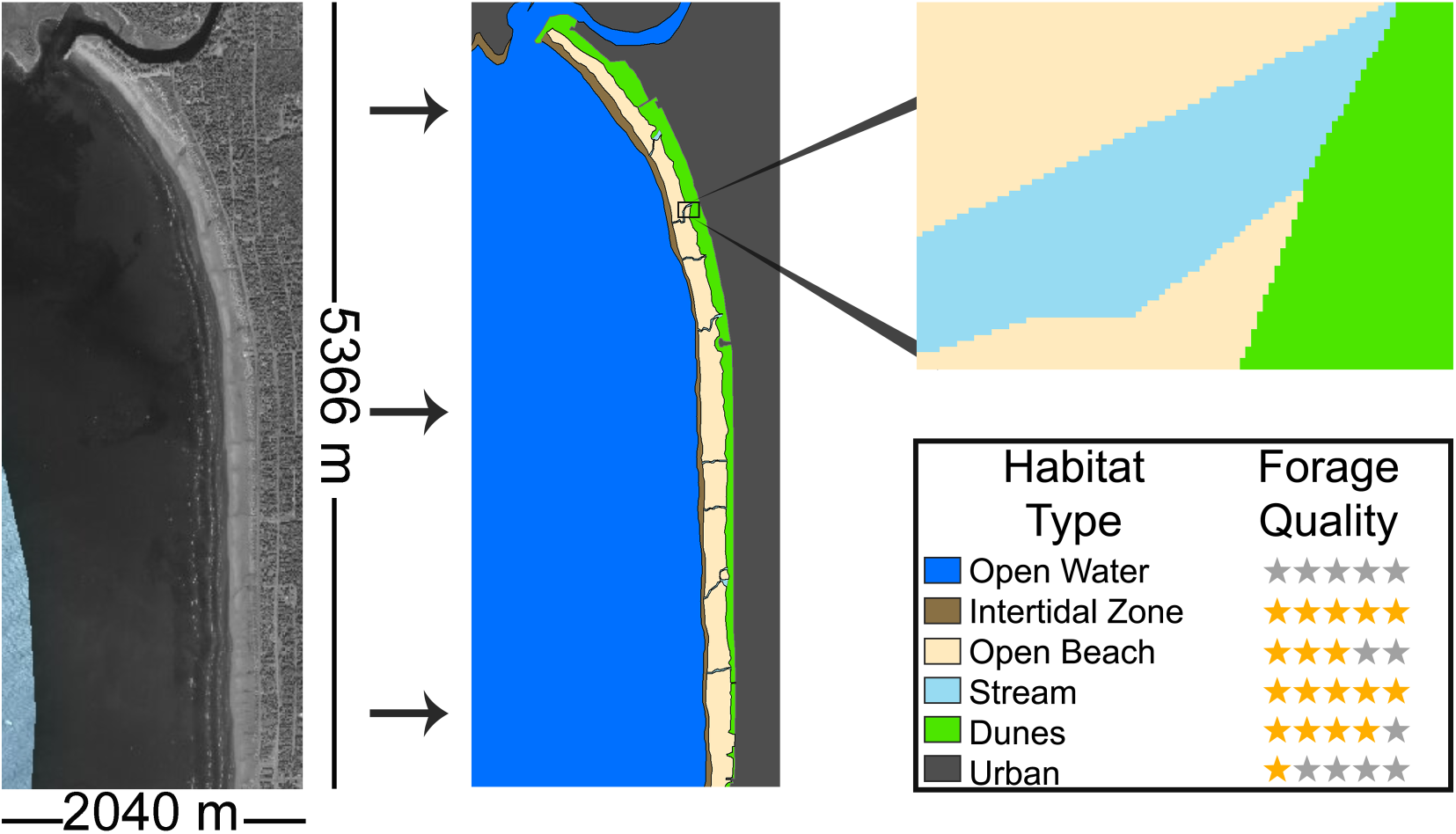
Digitization of Sauble Beach, Ontario. An aerial photograph comprised of a 5366 m long by 2040 m wide area of Sauble Beach was imported into ArcMap where regions could be coloured based on their habitat type. Each habitat type has an associated foraging quality that determines the amount of energy that a piping plover can gain in that habitat, with open water being the worst and intertidal and stream being the best.

Since enviro-ABMs treat partitioned environment cells as agents, the partitions of the physical layer become the agent units in a given simulation. Each agent stores abiotic information about its corresponding 1 m × 1 m plot of land; in this case, each agent stores what habitat type is contained in the given plot. By themselves, the agents do not have much functionality; the biological layer and anthropogenic layers are what provide the functionality to the agents and to the model itself.

### 2.3 Biological Layer

The biological layer for this seeks to simulate processes related to the distribution, abundance, systematics, behaviour, and energetics of Piping Plovers at Sauble Beach. Our study only focused on the growth of Piping Plover chicks within the first month, and so only functionality related to nesting and foraging (including subsequent movement and energy gains) were implemented.

#### 2.3.1 Nesting

At the beginning of a simulation, nests are randomly injected into the model. The model cycled through all the agents that were the “sandy beach” habitat (corresponding with Great Lakes Piping Plovers’ nesting habits of excavating a “scrape” to deposit their eggs), and a random Bernoulli trial (*P* (*Success*) = 1.00×10*^−^*^5^) was conducted for each agent. A nest was created if a 1 was drawn; if a 0 was drawn, the model moved to the next agent and a nest was attempted there using the same Bernoulli trial. Once a nest is placed, a rule is created that no other nest can be placed within a 200 m radius of that nest (Cairns 1982). The abundance of nests depends on the number of adults in the simulation: the model assumes a 50/50 split of male and female adults, and so the abundance of nests will be half of the number of adults. The abundance of initial adults at the beginning of the simulation is chosen by the user before running the model. For our study, the adults do not serve any purpose other than as a representation for breeding potential. Additionally, the nests themselves do not provide much functionality other than as a starting location for clutches of Piping Plover hatchlings and for a “home location” for Piping Plover chicks to go to at the end of the day.

#### 2.3.2 Foraging and Energy Gain

In the model, a single Piping Plover chick is represented as a decimal number corresponding to its mass at a given time. For simplicity, we kept clutches of Piping Plovers together through time, so the biological unit to be stored in a given agent is simply a vector of Piping Plover weights with the length of the vector corresponding to the size of the clutch. Additionally, we kept a constant clutch size of 4 as this is the typical clutch size for Piping Plovers (Cairns 1982).

Initial Piping Plover chick weight is drawn from a normal distribution with *µ* = 6 grams and *σ*^2^ = 0.5 (Wilcox 1959, Cairns 1982, Elliott-Smith and Haig 2004). Piping Plovers gain energy by foraging in the agent they are in. Different habitat types have different foraging quality, and so Piping Plovers will gain more energy in agents with higher foraging quality. With that, Piping Plover movement follows a random walk. To determine where a Piping Plover will move to, the following steps are taken. For a given active agent containing a Piping Plover:

1. Create a matrix *A* of all agents within a 25 m radius of the active agent’s location (Cohen 2014)
2. Map each agent in *A* with its corresponding foraging quality (given the habitat type contained in the agent) to create a new matrix *B*.
3. Normalize matrix *B*; now we have a probability matrix *P* such that each entry is the probability that the Piping Plover moves to that agent.
4. Use the probability matrix *P* to draw a random sample from a multinomial distribution; the index of the first success (indicated by a 1) will be where the Piping Plover moves to.

### 2.4 Anthropogenic Layer

The anthropogenic layer for this model can be thought of as the experimental treatment being applied to either the physical layer, biological layer, or both. In our study, we experimented with two anthropogenic layers: a human presence layer – that is, for each agent, there is a probability that there is human presence at that location; and a human exclosure layer – that is, a rule that humans cannot go within an *r* metre radius of a nest.

Implementation of these layers are discussed later. However, we will note here how these anthropogenic layers affect the other layers. Specifically, our study has these anthropogenic layers affecting the biological layer, particularly the random walk Piping Plovers take when foraging. When there are humans in nearby foraging areas, Piping Plovers will not move to that area to feed, regardless as to whether it may provide better foraging. With this, we can update our random walk for Piping Plover foraging:

1. Create a matrix *A* of all agents within a 25 m radius of the active agent’s location

a. Anthropogenic layer effect: remove all entries in *A* that contain humans or predators so that Piping Plovers do not move there. This creates an updated matrix *_A_′*
2. Map each agent in *A^′^* with its corresponding foraging quality (given the habitat type contained in the agent) to create a new matrix *B*.
3. Normalize matrix *B*; now we have a probability matrix *P* such that each entry is the probability that the Piping Plover moves to that agent.
4. Use the probability matrix *P* to draw a random sample from a multinomial distribution; the index of the first success (indicated by a 1) will be where the Piping Plover moves to.

This random walk is summarized in Figure 2.

**Figure 2:**
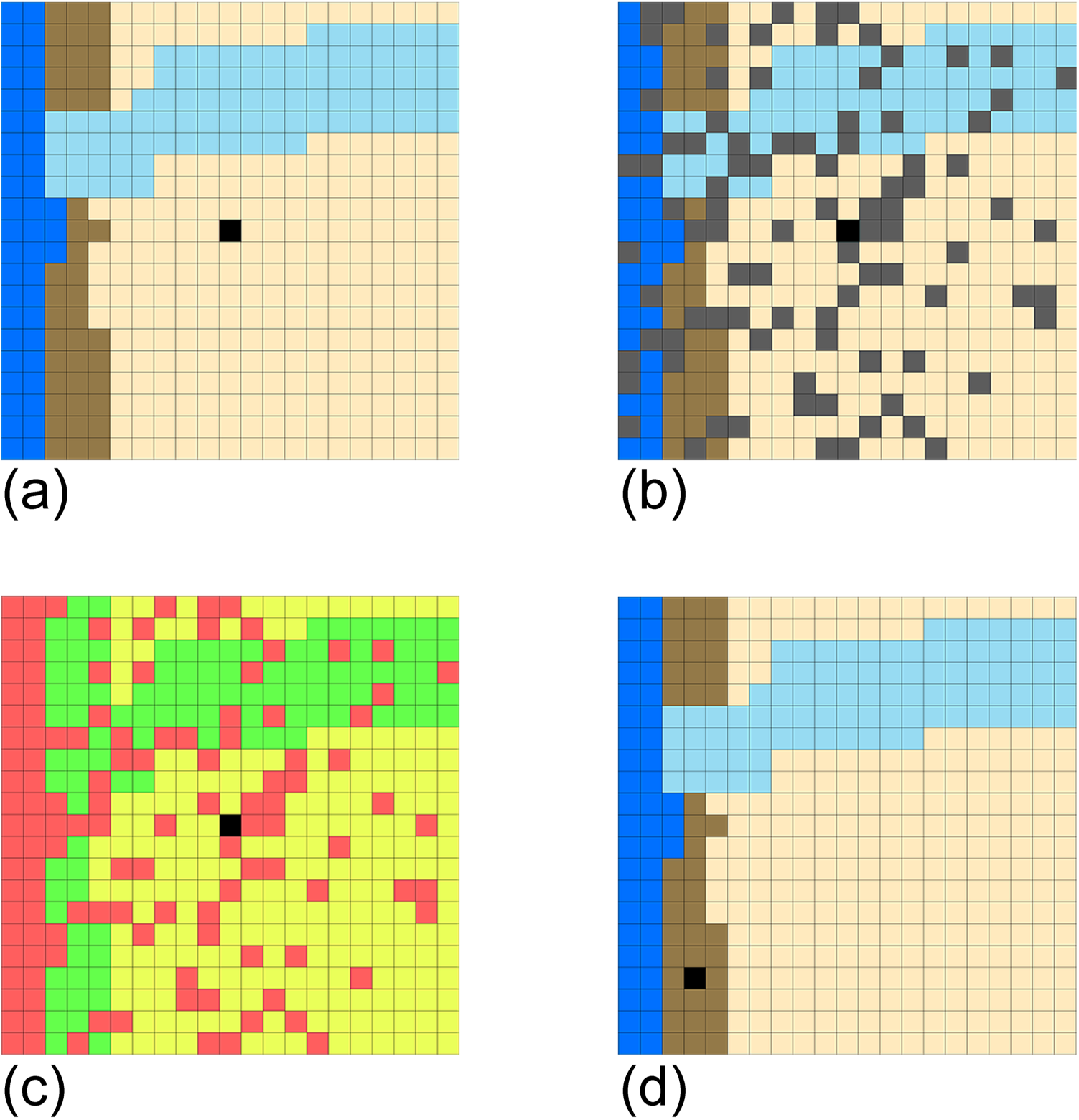
Random walk of piping plovers given surrounding habitat type and human presence. For simplicity, this figure shows a 10 m wide radius of potential agents (a) to which a piping plover (black square) could move. Any agents that contain humans or predators are removed as options (b), and the remaining options are mapped to probabilities based on their habitat type (c). A green square in (c) would imply a higher probability to move there than a yellow or red. These probabilities are used in a multinomial distribution where a random sample is drawn and mapped back to a new location where the plover moves to (d).

Though the Piping Plover will always move to an agent that does not contain humans, there could still be humans nearby. Once at a foraging location, the Piping Plover checks if there are humans in its “alert distance” of 50 m (Gulbransen et al. 2006, Cohen 2014). If humans are nearby, the Piping Plover forages at a reduced rate, simulating a real-life Piping Plover exhibiting on-alert behaviours such as crouching rather than spending its time feeding (Burger 1992). If no humans are nearby, then the Piping Plover forages normally, gaining energy at the typical rate.

### 2.5 Process Overview and Scheduling

The model proceeds in 5-minute time steps. For this study, chick weight was used as a response variable, and because Piping Plovers are most vulnerable in their first month of life (that is, after hatching but prior to fledging), a 31-day time period was used (Elliott-Smith and Haig, 2004).

For each time step, the model iterates through a list of active agents; that is, agents that contain at least one Piping Plover. A decision tree is then used for each active agent. If there is human presence in the agent, any Piping Plover in the agent would simply flee. If there is no human presence in the agent, the model checks if it is time to forage (simply determined by whether it was “day” or “night”). If it is time to forage, the model checks if there are humans in the alert distance of 50 m as discussed in Foraging and Energy Gain. If there are humans nearby, then the Piping Plover forages at a reduced rate of 50% its normal foraging. Otherwise, if there are no humans nearby, it forages as normal. The energy gain per time step *t* given no humans in alert distance is described by

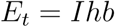

where *I* is an indicator function of whether humans are in alert distance (*I* = 1) or not (*I* = 0.5) *h* is an energy multiplier that corresponds to the current habitat type of the Piping Plover (Table 2; Cohen 2014), and *b* is a base amount of energy gained per time step in grams. This base amount of energy is given by

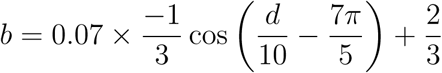

where *d* is the numerical day. This base energy gain per time step is simply a calibration parameter to ensure the Piping Plover growth rate reasonably matches the growth rate found by Martens and Goossen (2008). It is dependent on the day *d* and is a sinusoidal curve to account for the change in concavity of the growth rate, discussed in Analysis.

**Table 2:** Energy multipliers by habitat type. Higher quality habitat type such as intertidal zone or streams will correspond with a higher energy gain per time step.

| Habitat Type | Energy Multiplier |
| --- | --- |
| Open Water | 0.00 |
| Intertidal Zone | 1.00 |
| Open Beach | 0.50 |
| Stream | 1.00 |
| Dunes | 0.75 |
| Urban | 0.00 |

Finally, if it is not time to feed (i.e. it is night time and it is time to rest), the model checks if the Piping Plovers are currently at their nest. If they are not, Piping Plovers move toward their nest. Otherwise, they simply rest in the agent they reside in; that is, they do not gain nor lose energy, and no anthropogenic disturbance occurs at night. Although there is some evidence to show Piping Plovers forage regularly at night, a study by Staine and Burger (1994) showed that nighttime foraging took place at a substantially lower rate than daytime foraging, and the intensity of nighttime foraging was highly variable. For this reason, we opted to not have simulated Piping Plovers forage at night. A summary of the decision tree is found in Figure 3.

**Figure 3:**
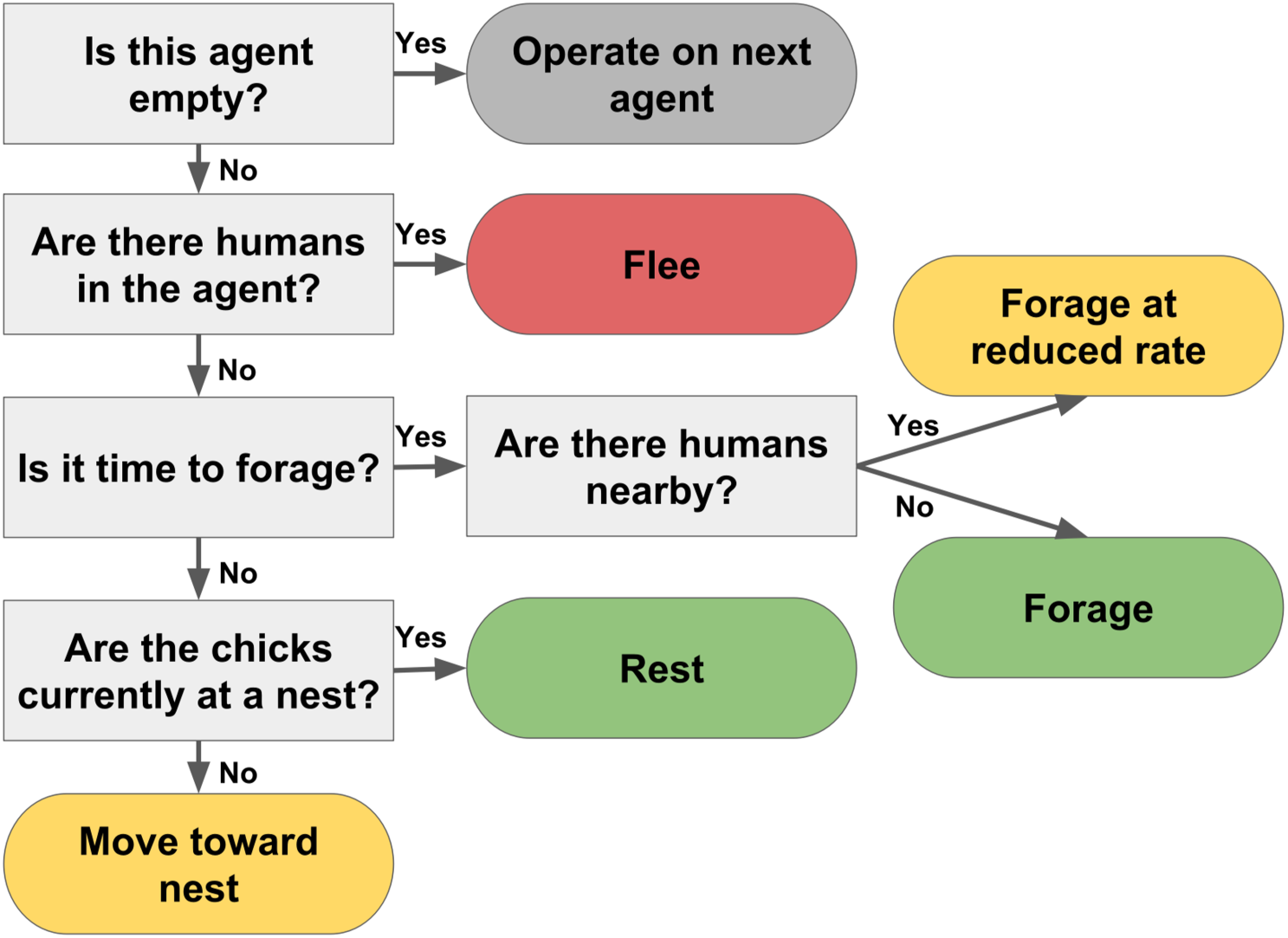
Flow chart of a typical simulation acting on a single enviro-agent.

### 2.6 Base Model

Using the previously described model, we first developed a base model. A base model is used as a “perfect world” scenario to compare experimental treatments of anthropogenic effects. In our study, our base model was simply a simulation where simulated Piping Plovers were free to forage without disturbance of humans. The goal was to model this after the Piping Plover growth measured at Chaplin Lake, SK (Martens & Goossen 2008). Chaplin Lake is designated as an Important Bird Area and experiences very little human interaction; therefore, Piping Plovers chicks at Chaplin Lake gain energy in a more ideal way as they can choose the best foraging locations to feed rather than choosing a lesser foraging location only because it does not contain humans. In total, we ran 50 simulations for the base model. Using the base model as baseline, anthropogenic layers can be incorporated into the model and Piping Plover growth can be compared. As mentioned, the anthropogenic layers of an enviro-ABM generally serve as the experimental treatment to the model. Thus, two anthropogenic layers were developed to be tested with this enviro-ABM implementation: (1) A layer simulating human presence and (2) A layer containing simulated human exclosures.

### 2.7 Experiment 1: Human Presence Layer

Human presence was simulated based on probabilities of humans occurring in each agent. We tested three levels of human presence: 20%, 40%, 60%; that is, at any given time step, any agent has either a 20%, 40%, or 60% chance of containing human presence. In this study, we did not classify type of human per agent, nor did we quantify the number of humans per agent; only a Boolean value was used to denote whether there was human presence in an agent. In total, we ran 20 simulations per level.

### 2.8 Experiment 2: Human Exclosure

The second anthropogenic layer was the addition of human exclosures. Human exclosures created a rule such that, given a radius *r*, no humans could be within *r* metres of a nest location. This was true no matter what level of human presence was being used for a simulation. This allowed Piping Plover chicks a “buffer zone” to forage without the disturbance of humans. Indeed, Piping Plover chicks can move in or out of this buffer zone as the opportunity arises (e.g. when undisturbed optimal foraging habitat occurs outside of this buffer zone). In this study, we experimented with the use of 100 m human exclosures; that is, all agents within 100 m of a nest could not contain humans. In total, we ran 20 simulations per each level of human presence considered in Experiment 1.

### 2.9 Analysis

For each of the base model, experiment 1, and experiment 2, we fitted simulated growth data to the following model equation:

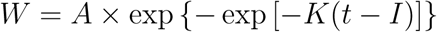

where *A* is the asymptotic mass (i.e. adult Piping Plover mass), *I* is the inflection point in the growth curve (where the rate of change of growth goes from increasing to decreasing), *K* is a growth rate constant, and *W* is the mass of the Piping Plover after *t* days. This particular parameterization of the Gompertz model was used by Martens and Goossen (2008) to fit their growth data, and is further discussed by Tjørve and Tjørve (2017) in their paper comparing various parameterizations. The growth data were fitted using the easynls package in R (Arnhold 2017). For this study, we are only concerned with comparing estimates of the asymptotic mass *A* and the growth rate *K*. Parameter estimates from these model runs were compared using Welch Two Sample t-test to estimates obtained from a study conducted by Martens and Goossen (2008) in Chaplin Lake, SK.

In experiment 1, 20 simulations were run for each level of human presence. The growth data for experiment were fitted using easynls to equation 1. The asymptotic mass and growth rate were compared to those from the base model using Welch Two Sample t-test.

In experiment 2, after introducing the use of the 100 m human exclosure, 20 simulations were run for each level of human presence as listed in experiment 1, and growth data were fitted to equation 1. The asymptotic mass and growth constant were compared to those from the base model using Welch Two Sample t-test.

Significance for both experiments was assessed at *α* = 0.05.

All analyses were completed using R: A Language and Environment for Statistical Computing (R Core Team 2017). Simulations and analyses were run on a Toshiba Satellite S855D with a 2.3GHz AMD Comal Quad-Core Mobile Processor A10-4600M and 8GB 1600MHz DDR3 RAM.

## 3 Results

### 3.1 Base Model Estimates

The mean chick weight across 50 simulations after 31 days was 44.72 g. The mean estimates of A, I, and K were 57.55, 2.390, and 0.072, respectively, with standard deviations of 2.242, 0.038, and 0.002, respectively. Relative bias was used as an indicator of how much the model may have overestimated or underestimated these parameters compared to parameters obtained by Martens and Goossen (2008). The median relative bias of A, I, and K were 8.469%, -78.77%, and -13.73%, respectively (Table 3). Figure 4 compares the target growth curve to the mean growth curve of 50 base model simulations.

**Figure 4:**
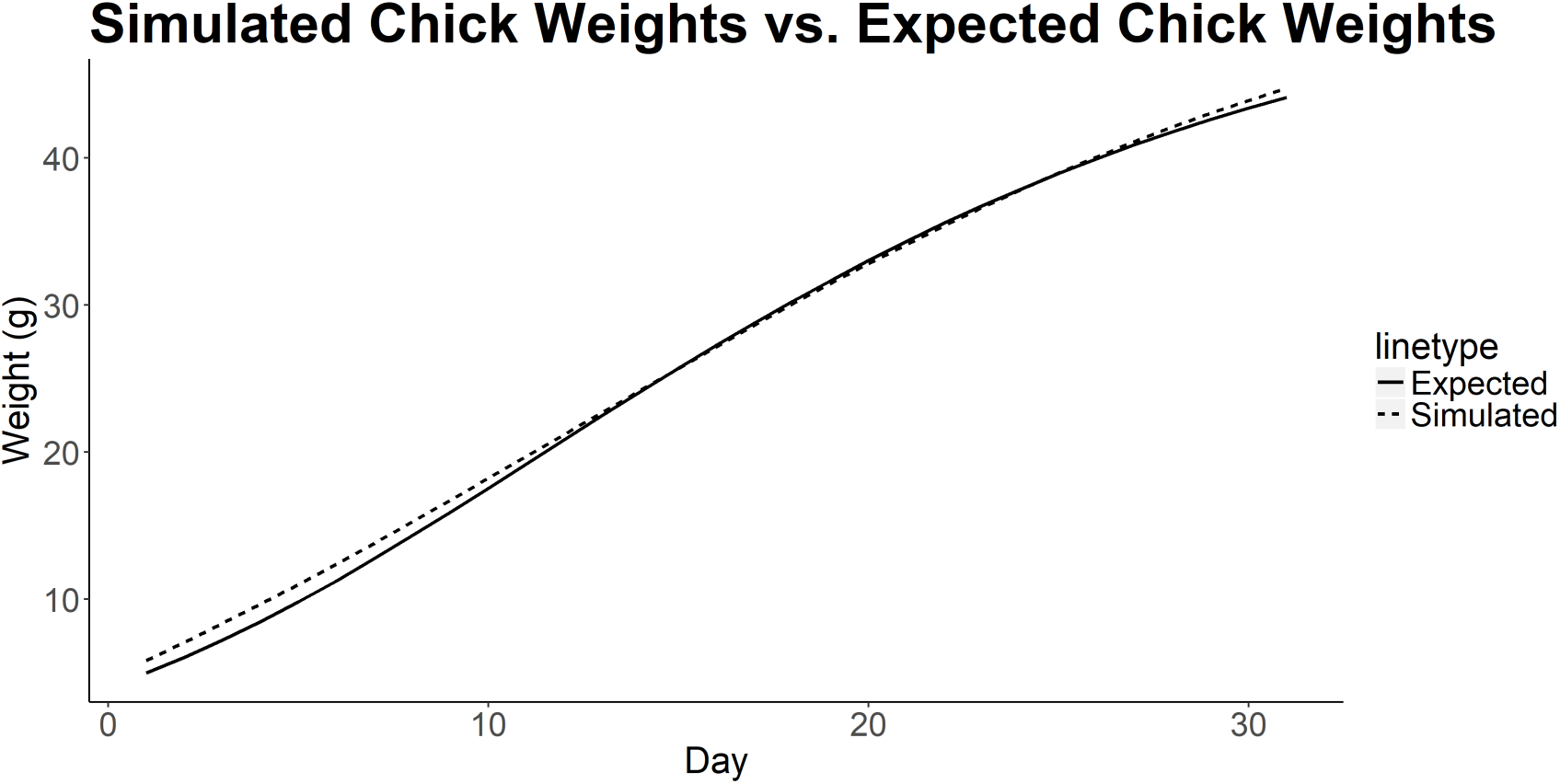
Time series plot of multiple base model runs (N = 50) simulating growth rate of piping plovers in an environment with no anthropogenic effects. Solid curve shows expected growth rate. Dashed curve shows the mean simulated weight per day.

**Table 3:** Parameter estimates of the adult weight in grams *A* and growth rate *K* from *N* =50 base model simulations, along with their standard deviation and median relative bias.

| Parameter | Fitted Estimate | Standard Deviation | Median Relative Bias |
| --- | --- | --- | --- |
| $A$ | 57.55 | 2.242 | 8.469% |
| $K$ | 0.072 | 0.002 | -13.73% |

### 3.2 Experiment 1

The mean chick weights after 31 days for the 20%, 40%, and 60% levels of anthropogenic activities (20 simulations each) were 9.77%, 17.24%, and 26.72% less than the base model with mean 31-day chick weights of 40.35 g, 37.01 g, and 32.77 g, respectively.

At the 20% level of human presence, the mean asymptotic mass was estimated at 52.17 g, a significant drop from the base model (*p <* 0.001). The mean growth rate estimate was calculated to be 0.071, which was also a significant drop in growth rate from the base model (*p <* 0.001).

The 40% level of human presence had even smaller values than the base model, with a mean asymptotic mass estimate of 48.33 g (*p <* 0.001) and a mean growth rate estimate of 0.069 (*p <* 0.001).

The trend continues at the 60% level of human presence, with a mean asymptotic mass estimate of 42.99 g (*p <* 0.001) and a mean growth rate estimate of 0.067 (*p <* 0.001).

Table 4 summarizes these parameter estimates for experiment 1.

**Table 4:**
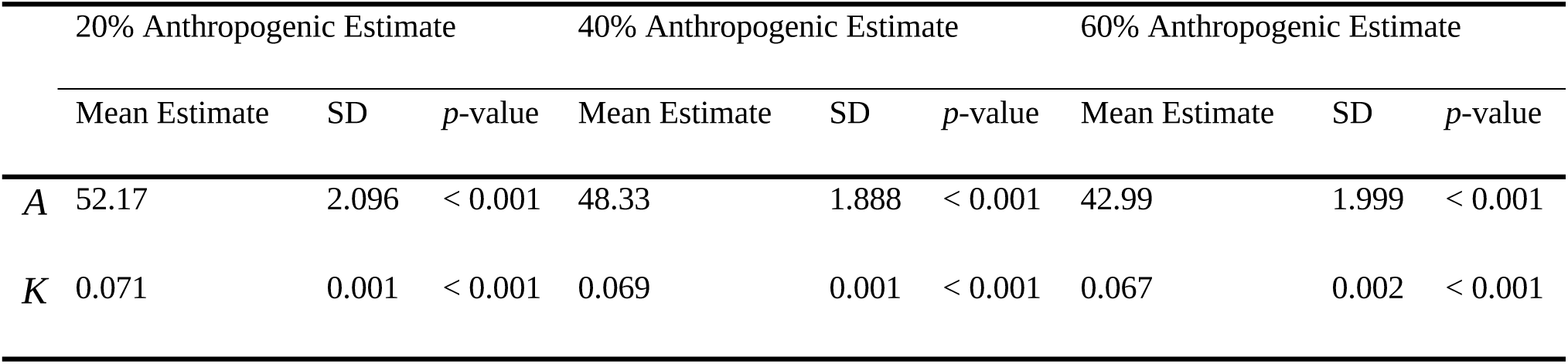
Results of Experiment 1. Experiment 1 tested the effects of increased human presence on the growth of Piping Plover chicks. Parameter estimates for adult weight in grams *A* and growth rate *K* were obtained from 20 simulations of each level, as well as their standard deviation and a p-value indicating their difference from the base model runs shown in Table 2.

### 3.3 Experiment 2

The mean chick weights after 31 days for the 20%, 40%, and 60% levels of anthropogenic activities with 100 m human exclosures (20 simulations each) were 2.26%, 5.01%, and 8.25% less than the base model, with mean chick weights after 31 days of 43.71 g, 42.48 g, and 41.03 g, respectively.

At the 20% level of human presence with a 100m exclosure, the mean estimate of asymptotic weight was 56.23 g, which was still a significant drop from the base model (*p* = 0.038). The mean growth rate estimation was calculated to be 0.0722, which was not significantly different from the base model (*p* = 0.594).

At the 40% level of human presence with a 100m exclosure, the mean asymptotic mass was estimated at 54.78 g, a significant drop from the base model (*p <* 0.001). The mean growth rate estimation was calculated to be 0.716 which was not a significant change from the base model (*p* = 0.560).

Finally, at the 60% level of human presence with a 100m exclosure, the mean asymptotic mass was estimated at 53.19 g which was significantly less than the base model (*p <* 0.001).

The mean growth rate estimate was calculated to be 0.0708, which was also significantly less than the base model (*p <* 0.001).

Table 5 summarizes these parameter estimates for experiment 2.

**Table 5:** Results of Experiment 2. Experiment 2 tested the effects of a 100 m exclosure on the growth Piping Plover chicks given the three levels of anthropogenic presence. Parameter estimates for adult weight *A* and growth rate *K* were obtained from 20 simulations of each level, as well as their standard deviation and a p-value indicating their difference from the base model runs shown in Table 2.

|  | 20% Anthropogenic Estimate |  |  | 40% Anthropogenic Estimate |  |  | 60% Anthropogenic Estimate |  |  |
| --- | --- | --- | --- | --- | --- | --- | --- | --- | --- |
| | Mean Estimate | SD | $p$-value | Mean Estimate | SD | $p$-value | Mean Estimate | SD | $p$-value |
| $A$ | 56.23 | 2.337 | 0.038 | 54.78 | 1.134 | < 0.001 | 53.19 | 1.651 | < 0.001 |
| $K$ | 0.072 | 0.002 | 0.594 | 0.072 | 0.002 | 0.056 | 0.071 | 0.001 | < 0.001 |

## 4 Discussion

The goal of this paper was to explore the use of enviro-ABMs in risk assessment of Piping Plovers breeding at Sauble Beach. We developed and described an enviro-ABM for simulating a population of Piping Plover chicks, we investigated a set of input parameters to create a base model (i.e. a “perfect world” scenario in which there are no anthropogenic disturbances to nesting Piping Plovers), and we developed and tested a set of scenarios to simulate anthropogenic activity for which to compare to the base model.

### 4.1 Parameter Estimates in the Base Model

Our base model gave reasonable simulations for Piping Plover growth given no anthropogenic disturbance. After running 50 simulations, the mean 31-day growth curve of Piping Plover chicks closely followed the growth curve found by Martens and Goossen (2008). This close match allows us to provide a baseline to which future experiments (i.e. human presence and human exclosures) can be compared. That is, we, as the model developers, provided the calibrating parameters to the model in order to ensure a good fit to use for further experimentation. Our statistical tests comparing our simulations to the real-world growth data serve to provide evidence that our base model growth rate is insignificantly different from data that we identified as an appropriate base model.

The base model tended to slightly overestimate the asymptotic mass, with a median relative bias of 8.47%. The mean asymptotic mass estimate of 57.55 g implies that once the simulated Piping Plovers reach their adult stages, their mass will fall around 57.55 g. Although this asymptotic mass is significantly greater than the asymptotic mass of 53.4 g used by Martens and Goossen (2008), 57.55 g still falls within the adult weight range of 43.0 – 63.0 g (Cairns 1982). When observing the growth rate estimates, the model tended to slightly underestimate the growth rate, with a median relative bias of -13.73%.

The inflection point of the growth curve was also severely underestimated. The inflection point is the point on the growth curve where the concavity changes; in the case of Piping Plover growth, it is the point where the rate of change of growth goes from increasing to decreasing, signifying that the plovers are still growing but are slowing down in their growing (typically when they are approaching their adult weight). This did not have any effect on our study as we were mainly concerned with the overall trend that the anthropogenic stressors might add to the model; our study was not focused on the day-to-day Piping Plover weights leading up to the day 31 weights.

It should be noted that the parameter estimates obtained from Martens and Goossen (2008) were determined using empirical data obtained from Chaplin Lake, SK; they are not necessarily indicative of true asymptotic mass or growth rates of the Sauble Beach Piping Plovers. Therefore, though the asymptotic mass tended to be slightly overestimated and the growth rate tended to be slightly underestimated, we were still comfortable with using these parameters as a base model that provided reasonable simulations of Piping Plover growth, and any experimentation done could accurately show a trend (if one exists) in Piping Plover growth.

### 4.2 Effects of Human Presence (Experiment 1)

When adding in human presence, the simulated Piping Plover chick weight tended to have lower estimated growth rates and lower asymptotic mass compared to the base model. These estimates decreased as the probability of human presence increased. In beach habitats with more humans, the Piping Plover chicks tend to spend more time on alert or fleeing from humans rather than spending time foraging, leading to the smaller growth rate (Cohen 2014). That said, both the 20% and 40% level of human presence had asymptotic mass estimates still within the normal range of adult Piping Plover masses of 43.0 – 63.0 g (Cairns 1982), so it is certainly possible for the Piping Plovers to hit normal adult mass despite the presence of humans, which is consistent with real-world observations. In observing chick weight after 31 days, the lowest amount of anthropogenic presence we investigated still showed close to a 10% drop in mean chick weight, with that percent drop increasing close to 28% at the highest amount of anthropogenic presence. These results show a clear trend in a decrease in Piping Plover growth as more anthropogenic activity is added to the environment. These results are consistent with weights of piping plovers on Sauble Beach as measured during the banding process (Hamm 2016), as well as with other studies that have looked at the effects of anthropogenic disturbances on other shorebird species (see Ruhlen et al. 2003, Yasúe & Dearden 2006, Liley & Sutherland 2007, Weston & Elgar 2007).

### 4.3 Effects of Human Exclosures (Experiment 2)

When adding a human exclosure, the asymptotic mass and growth rate were still lower than the base estimates, but certainly less so. Although the Piping Plovers still had a lower growth rate and lower asymptotic mass than the base model, the estimates were higher than estimates without human exclosures. This would imply that the use of human exclosures may help with mitigating the effects of anthropogenic presence. In comparing the chick weights after a 31 day simulation from experiment 1 and experiment 2, we see that the use of human exclosures tended to reduce the decrease in weight as more human presence was added to the simulation (Figure 5). This is especially true in looking at the decrease in mean weight after 31 days, as the percent drop did not surpass 10% at the highest level of anthropogenic presence.

**Figure 5:**
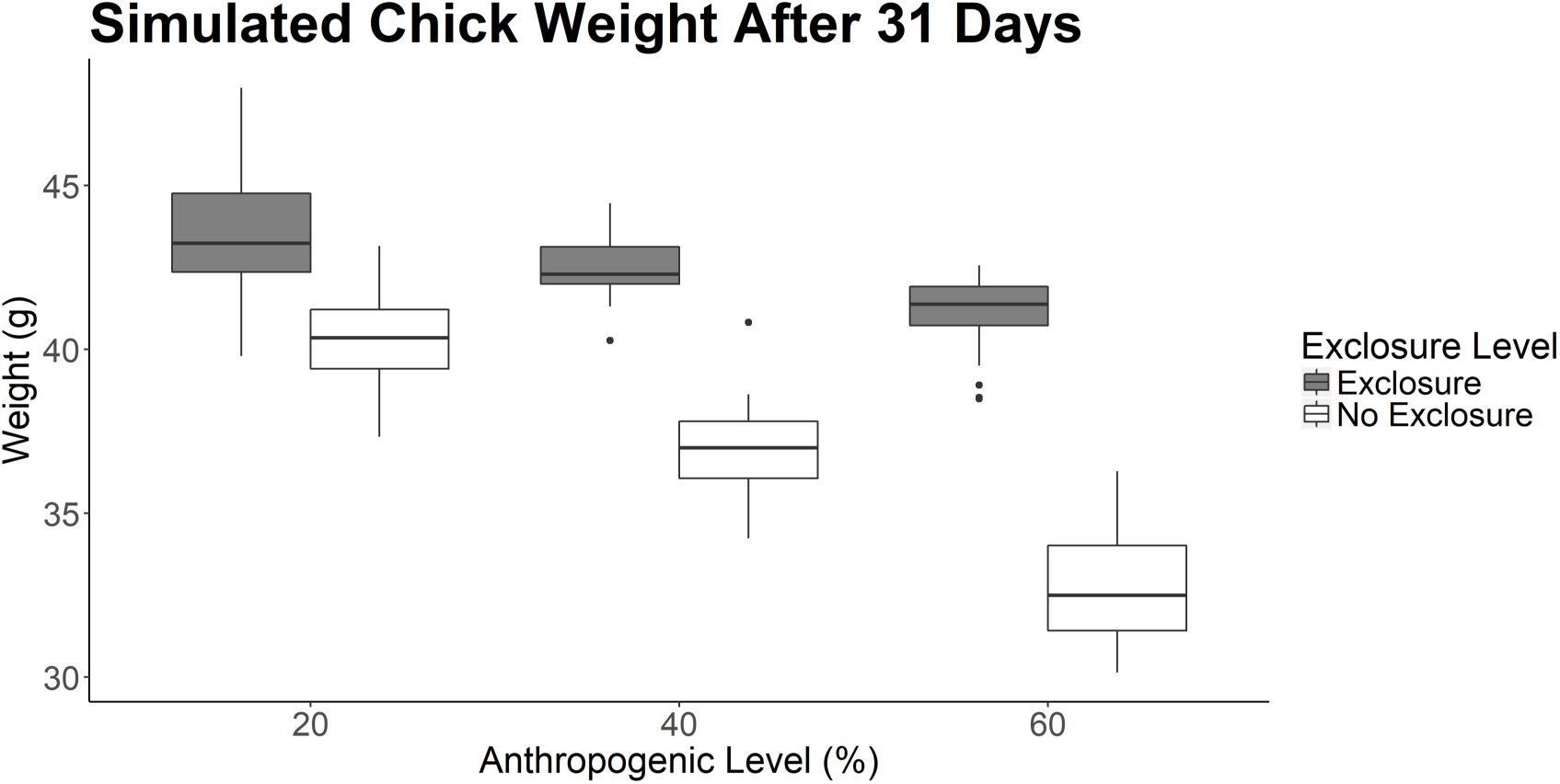
Boxplot of simulated chick weights after a 31-day simulation grouped by anthropogenic presence level and separated by whether an exclosure was used.

In our model, the use of a human exclosure significantly increased the Piping Plovers’ growth rate back to a level that was close to our base model, despite an increase of anthropogenic presence. Human exclosures are a common technique used in Piping Plover management as it allows for a barrier to decrease human and predator interactions with nesting adult plovers and vulnerable hatchlings (Johnson & Oring 2002). We predicted that if exclosures provide a buffer zone of piping plovers to feed uninhibited by human presence, then their growth rate should increase slightly toward the ideal growth rate. Our study reaffirms that the use of human exclosures may mitigate the effects of human presence on growing Piping Plover chicks. Further field studies should look to compare the growth rate of piping plovers with the use of an exclosure compared to those without the use. Additionally, further studies using this model can experiment with varying sizes of human exclosures to investigate a minimum exclosure size that can successfully mitigate the effects of human presence, or with different shapes of the exclosure, such as one that extends toward the shoreline as suggested by Hamm (2016).

### 4.4 Future Work

The creation of any type of simulation model, including enviro-ABMs, rely heavily on information that is already known about the species of interest and its environment. An issue that we faced for this study was the limited available data on ideal growth curve of Piping Plovers at Sauble Beach. We resorted to using a growth curved obtained from Chaplin Lake which contains very little anthropogenic activity (Martens & Goossen 2008). One major goal in further developing this model is to reach out to biologists and government officials to develop a plan to monitor growth rates of Piping Plovers at Sauble Beach and in the general Great Lakes region. There exists a small amount of mass measurements when chicks are measured by Canadian Wildlife Service officials upon banding (Hamm 2016), but to our knowledge there does not exist a “full” range of mass measurements like those described by Martens and Goossen (2008).

Simulating the foraging behaviour of Piping Plovers relies heavily on information that is known about their feeding habits. To improve our estimates of energy gain through time, we recommend that future field studies should be done to investigate Piping Plover feeding habits, especially as it relates to invertebrate abundance on beaches. Our model currently makes assumptions of habitat quality (and therefore foraging quality) based on some preliminary work from Hamm (2016), habitat selection results from Le Fer et al. (2008) and Haffner et al (2009). and modelling work of piping plovers from Cohen (2014), and the random walk and energy gains of the simulated piping plovers in this model are based heavily on these assumptions. With that in mind, if the habitat quality of Sauble Beach (or some other site that a researcher may want to simulate) were to differ greatly from these assumptions, our model may no longer be as accurate as it could be. As such, we recommend that researchers looking to adapt this model for a different environment should pay close mind to the habitat quality of the site in mind. This is easily accomplished when digitizing the map of the site as the researcher can simply create their own habitat types, but it should not be taken for granted that the habitat types and foraging qualities will be the same across environments.

We recommend that implementations of this model should seek to develop a way to simulate predation, to both Piping Plover eggs and Piping Plover chicks. This should be combined with field studies investigating the growth rate of Piping Plovers on a human-use beach as well as field studies investigating Piping Plover feeding habits. As mentioned, unfledged Piping Plovers are the most vulnerable to predation. An issue seen at Sauble Beach is the increase of human-subsidized predators such as gulls and weasels, so a model that implements predation along with new information on feeding habits and response to humans would be of great value in assisting with management policy of Piping Plovers at Sauble Beach (Hamm 2016, Stranger 2017).

Given continual studies into the life history of Piping Plovers, especially their breeding ecology and their use of microhabitats on their breeding grounds, this model should be used as a tool to inform future conservation and management efforts of the Piping Plovers. We have demonstrated here that the model can provide reasonable estimates of Piping Plover chick growth that is unimpeded by human interaction, and we have demonstrated the effectiveness of the model to simulate human presence and human exclosures. Since enviro-ABMs take a layered approach to their development, government agencies and biologists can jointly work with this model to develop a plethora of anthropogenic layers (simulating their potential management strategies) to test on a simulated population of Piping Plovers. This would eliminate the need to play a “guess-and-check” game of management technique implementation; biologists and government officials could test their management technique ideas in a simulation and develop a better sense of what management technique might be best to use in practise. We hope to be able to work with lead Piping Plover researchers throughout North America to be able to develop these simulation scenarios, and to develop research teams into further identifying knowledge gaps that this model may be able to identify and fill.

As the number of variables to consider are added into this model, especially if we begin to work with lead researchers, we expect that this model implementation could become computationally intensive, despite the benefits laid out by Rose et al. (2017) of using an enviro-ABM. A model developed by Liley and Sutherland (2007) sought to use a game theory approach to predict anthropogenic effects on Ringed Plovers *Charadrius hiaticula*. Though their model setup differs from ours in that ours is a spatially explicit model, we can consider using their method of PCA for variable selection to best inform our model as information to inform the model increases. That is, given a variety of potential variables to use in the model, we can adopt a PCA method similar to Liley and Sutherland (2007) to decide which variables would best be used in a spatially explicit model such as ours.

Though we only developed an implementation for use with Piping Plovers, we have seen the use of enviro-ABMs in other species such as Lake Whitefish (Rose et al. 2017), and so we are confident that other implementations of this model can seek to simulate the life history of other species at risk to develop timely management practices.

## Acknowledgements

The authors would like to thank Alicia Fortin from Plover Lovers, as well as the numerous volunteers who monitor the nests at Sauble Beach each year. We would like to thank the reviewers who provided extremely valuable input to improve the paper. Funding in part for this project through the Taverner Award from the Society of Canadian Ornithologists. We acknowledge that the University of Guelph resides on the ancestral lands of the Attawandaron people and the treaty lands and territory of the Mississaugas of the Credit. We recognize the significance of the Dish with One Spoon Covenant to this land and offer our respect to our Anishinaabe, Haudenosaunee and Métis neighbours as we strive to strengthen our relationships with them.

